# Lgl resets Par complex membrane loading at mitotic exit to enable asymmetric neural stem cell division

**DOI:** 10.1101/2024.09.29.615680

**Authors:** Bryce LaFoya, Sarah E. Welch, Kenneth E. Prehoda

**Author notes:** contributed equally (Bryce LaFoya, Sarah E. Welch).

## Abstract

The Par complex regulates cell polarity in diverse animal cells**^1–4^**, but how its localization is restricted to a specific membrane domain remains unclear. We investigated how the tumor suppressor Lethal giant larvae (Lgl) polarizes the Par complex in *Drosophila* neural stem cells (NSCs or neuroblasts). In contrast to epithelial cells, where Lgl and the Par complex occupy mutually exclusive membrane domains, Lgl is cytoplasmic when the Par complex is apically polarized in NSCs**^5^**. Importantly, we found that Lgl’s key function is not in directly regulating metaphase Par polarity, but rather in removing the Par complex from the membrane at the end of mitosis, creating a "polarity reset" for the next cell cycle. Without this Lgl-mediated reset, we found that residual Par complex remains on the basal membrane during subsequent divisions, disrupting fate determinant polarization and proper asymmetric cell division. These findings reveal a novel mechanism of polarity regulation by Lgl and highlight the importance of the pre-polarized state in Par-mediated polarity.

## Introduction

Cortical polarity is a fundamental cellular characteristic that underlies diverse biological processes, including directional cell migration, tissue organization, and asymmetric cell division. The Par complex, consisting of Par-6 and atypical Protein Kinase C (aPKC), plays a pivotal role in establishing and maintaining polarity in a wide range of animal cells, from epithelial cells to neurons^1–3,6^. The proper localization and function of the Par complex are particularly critical in stem cells, where asymmetric distribution of cell fate determinants governs the delicate balance between self-renewal and differentiation. A fundamental question in cell polarity research is how the Par complex itself is restricted to specific membrane domains. The tumor suppressor Lethal giant larvae (Lgl) plays a key role in Par polarity by restricting the Par complex to the proper membrane domain^5,7–10^, but how Lgl controls Par complex localization remains an open question.

Par polarity is the result of a complex interplay of positive and negative regulators. While factors like Par-3 (Bazooka in *Drosophila*) and Cdc42 ensure that the Par complex is targeted to the membrane^11–13^, the negative regulation provided by Lgl restricts the complex to a specific membrane domain. Lgl is thought to perform this critical function by mutual exclusion: the Par complex and Lgl reciprocally control one another’s localization by binding to the membrane^3,5,7,14^. The mutual exclusion model is exemplified by interphase epithelial cells, where the Par complex is confined to the apical plasma membrane while Lgl occupies the basolateral domain (Fig. 1A). Loss of either component, Lgl or Par complex, allows the other to enter the incorrect domain^5,7,9^. How the Par complex excludes Lgl from the Par domain is well-understood^15–17^, but little is known about how Lgl regulates Par complex localization.

**Figure 1.**
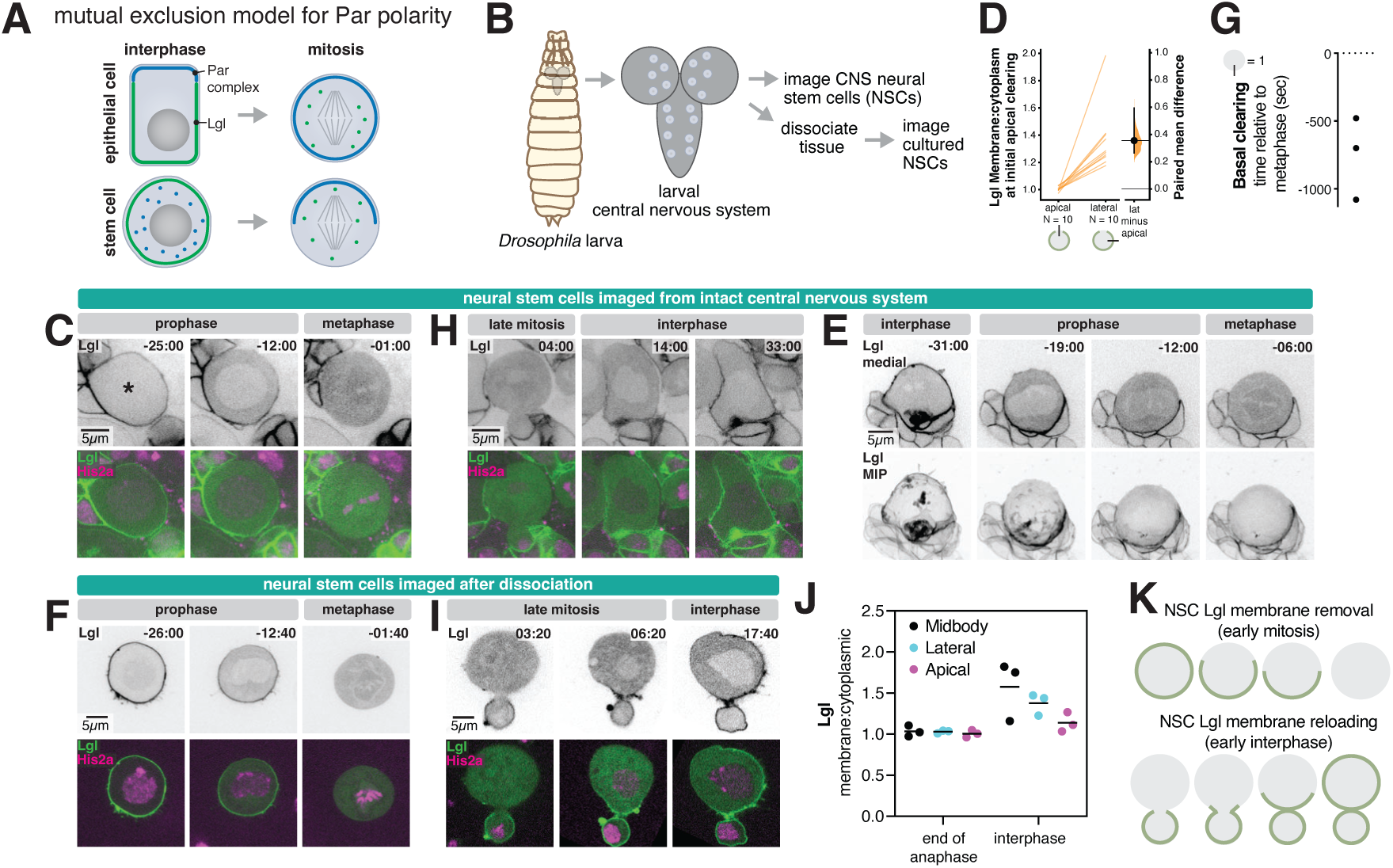
Patterned Lgl displacement and reloading during neural stem cell asymmetric division. (A) Mutual exclusion model for Par polarity. In this model, the Par complex (comprised of Par-6 and atypical Protein Kinase C or aPKC) establishes a mutually exclusive membrane domain with Lethal giant larvae (Lgl), wherein the presence of one protein on the membrane prevents the binding of the other. This model is consistent with the localization pattern in epithelial cells but not Drosophila neural stem cells (NSCs) and sensory organ precursors, where the Par complex becomes polarized during mitosis while Lgl remains cytoplasmic. (B) Imaging Drosophila larval brain neural stem cells (NSCs). The larval central nervous system (CNS), including the brain lobes and ventral nerve cord, was extracted from third-instar larvae (L3). Neural stem cells (NSCs), characterized by their relatively large size, specific locations within the brain, and expression of tissue-specific transgenes, were either imaged within the intact CNS or, alternatively, the tissue was dissociated to isolate and image the NSCs individually. (C) Removal of Lgl from the plasma membrane in early mitosis in an NSC from the intact CNS. Frames from Video 1 are shown. The NSC (marked with asterisk) is expressing Lgl-GFP (expressed from Worniu-GAL4 driven UAS) and Histone H2A (His2a-RFP). Time is shown relative to anaphase onset. (D) Patterned removal of Lgl from the plasma membrane in early mitosis. The ratio of membrane-to-cytoplasmic Lgl is presented for ten distinct neural stem cells (NSCs) in the apical and lateral regions during early mitosis, when Lgl was first cleared from the apical membrane. The Gardner-Altman estimation plot illustrates the paired measurements for each NSC. Additionally, the 95% confidence interval is displayed in the mean difference comparison, along with the corresponding bootstrap distribution from which it was derived. (E) A 3D projection of Lgl dynamics during early mitosis. Frames from Video 1 are shown. The NSC is expressing Lgl-GFP (expressed from Worniu-GAL4 driven UAS). The top row depicts a medial optical section. The bottom row depicts a maximum intensity projection (MIP) of optical sections across one hemisphere of the cell. Time is shown relative to anaphase onset. (F) Removal of Lgl from the plasma membrane in early mitosis in an NSC dissociated from the CNS. Frames from Video 1 are shown. The NSC is expressing Lgl-GFP (expressed from Worniu-GAL4 driven UAS) and Histone H2A (His2a-RFP). Time is shown relative to anaphase onset. (G) The timing of the clearance of the basal Lgl membrane signal, represented by a membrane-to-cytoplasmic signal ratio close to one, is presented for three dissociated neural stem cells (NSCs) in relation to metaphase. (H) Lgl reloading to the plasma membrane in early interphase in an NSC from an intact CNS. Frames from Video 1 are shown as in (C). Time is shown relative to anaphase onset. (I) Lgl reloading to the plasma membrane in early interphase in an NSC dissociated from the CNS. Frames from Video 1 are shown as in (F). (J) Lgl patterned membrane reloading in early interphase. The ratio of membrane-to-cytoplasmic Lgl is shown for two time points: one at the conclusion of anaphase and another during interphase. Measurements were taken near the cytokinetic midbody as well as along the lateral and apical membranes. (K) Schematic representation of Lgl membrane dynamics in neural stem cells (NSCs). During interphase, Lgl is distributed across the NSC membrane. In early mitosis, membrane clearing begins starting at the apical pole and fully clearing by prophase. Membrane reloading of Lgl begins in early interphase beginning near the midbody region.

Like epithelial cells, proper Par localization in NSCs requires Lgl^4,12,18,19^. In NSCs lacking Lgl function, the Par complex is localized across the entire membrane instead of being restricted to its proper domain^12,18^, similar to epithelia. However, a key difference between Lgl’s behavior in NSCs and epithelia lies in Lgl’s localization. Instead of localizing to a complementary membrane domain, Lgl is cytoplasmic when the Par complex is polarized in NSCs^5^ (Fig. 1A). Polarized Par complex with cytoplasmic Lgl has also been observed in sensory organ precursor (SOP) cells^20^, further challenging the exclusion model. Thus, Lgl’s behavior in NSCs and SOPs indicates that Lgl can regulate Par complex localization without being on the membrane.

## Results and Discussion

### Patterned Lgl membrane dynamics during NSC mitosis

To gain insight into how Lgl regulates Par complex localization in NSCs, we first examined Lgl’s membrane localization dynamics including how it is restored to the membrane following mitosis. We imaged Lgl’s localization in NSCs from *Drosophila* larval brain explants expressing Lgl-GFP (expressed by Worniu-GAL4 driven UAS) and Histone H2A-RFP (expressed from its endogenous locus) using rapid, super-resolution spinning disk confocal microscopy (Fig. 1B). Lgl removal from the membrane began in early prophase, initiating at a small area near the apical pole of the cell and spreading rapidly across the membrane such that it appeared to be completely removed by late prophase (Fig. 1C,D; Video 1). This pattern could be observed both in medial sections and maximum intensity projections of the full cell volume (Fig. 1E; Video 1). The full volume data also revealed how Lgl displacement from the membrane was affected by the membrane structures that are present on the NSC surface^21–23^. Lgl was enriched in the membrane structures, consistent with their higher membrane density. Lgl disappeared from the structures as it was displaced from the membrane, while the structures themselves remained evident when imaging a membrane sensor (Video 1).

While Lgl appeared to be completely removed from the membrane when imaging NSCs in intact tissue, progeny cells adhered to the basal region prevented a definite determination. We imaged individual cultured NSCs that had been dissociated from brains to observe basal Lgl dynamics in the absence of adhering cells. The Lgl dynamics in dissociated NSCs were consistent with our initial observations of apically-directed membrane clearing and provided unambiguous evidence that Lgl is indeed completely removed from the membrane by late prophase (Fig. 1F,G; Video 1). The patterned Lgl membrane displacement dynamics indicate that removal is a spatially regulated process, like that observed for SOPs^20^, rather than one in which it is simultaneously removed from the membrane.

NSCs undergo repeated cycles of asymmetric division such that Lgl must be restored to the membrane at some point to achieve its interphase localization state. We analyzed Lgl dynamics in both intact brain and dissociated NSCs to determine when and how Lgl is restored to the membrane following mitosis. Our results showed that Lgl becomes rapidly restored to the membrane at the completion of mitosis, beginning near the cytokinetic pore connecting the nascent sibling cells (Fig. 1H-J; Videos 1,3), similar to the dynamics observed in follicular epithelial cells^24^. Our results provide a detailed description of Lgl’s dynamic localization throughout the cell cycle in NSCs, revealing precise spatial and temporal control of its membrane association (Fig. 1K).

### aPKC removes Lgl from the NSC early in mitosis

Since Lgl is cytoplasmic at metaphase in NSCs, when the Par complex is polarized, we sought to understand how it transitions between membrane-bound and cytoplasmic states. Evidence from various cell types indicates that Lgl can be phosphorylated and removed from the membrane by the Par complex subunit atypical Protein Kinase C (aPKC) but also by the mitotic kinase Aurora A^5,7^. Aurora A could also regulate Lgl’s membrane association indirectly by influencing aPKC polarity^20,25^. We sought to determine whether one or both kinases are responsible for the patterned removal of Lgl from the NSC membrane during prophase. To dissect the relative contributions of aPKC and Aurora A to Lgl removal, we separately inhibited aPKC or Aurora A activity and assessed the effect on Lgl localization dynamics. We inhibited aPKC by expressing a UAS-controlled aPKC RNAi specifically in larval brain NSCs using Worniu-GAL4 driven expression. For Aurora A inhibition, we added the specific inhibitor MLN8237^7^ to the larval brain explant culture media.

In NSCs where aPKC was depleted by RNAi, we observed that Lgl removal from the membrane in mitosis was largely inhibited (Fig. 2A,B; Video 2). Unlike wild-type NSCs, where Lgl was removed from the membrane late in prophase, aPKC RNAi NSCs had persistent Lgl membrane enrichment throughout mitosis. We did detect a small decrease in apical Lgl membrane signal beginning in early prophase (Fig. 2A,B) suggesting the presence of residual aPKC activity or a small contribution from another mechanism. The retention of Lgl on the membrane in aPKC-depleted cells indicates that aPKC-mediated phosphorylation is the primary mechanism driving Lgl dissociation from the plasma membrane as NSCs enter mitosis. Interestingly, Lgl’s membrane enrichment during interphase was enhanced in aPKC RNAi NSCs (Fig. 2A,B), suggesting that Lgl exchanges between the membrane and cytoplasm due to phosphorylation and dephosphorylation in wild-type NSCs.

**Figure 2.**
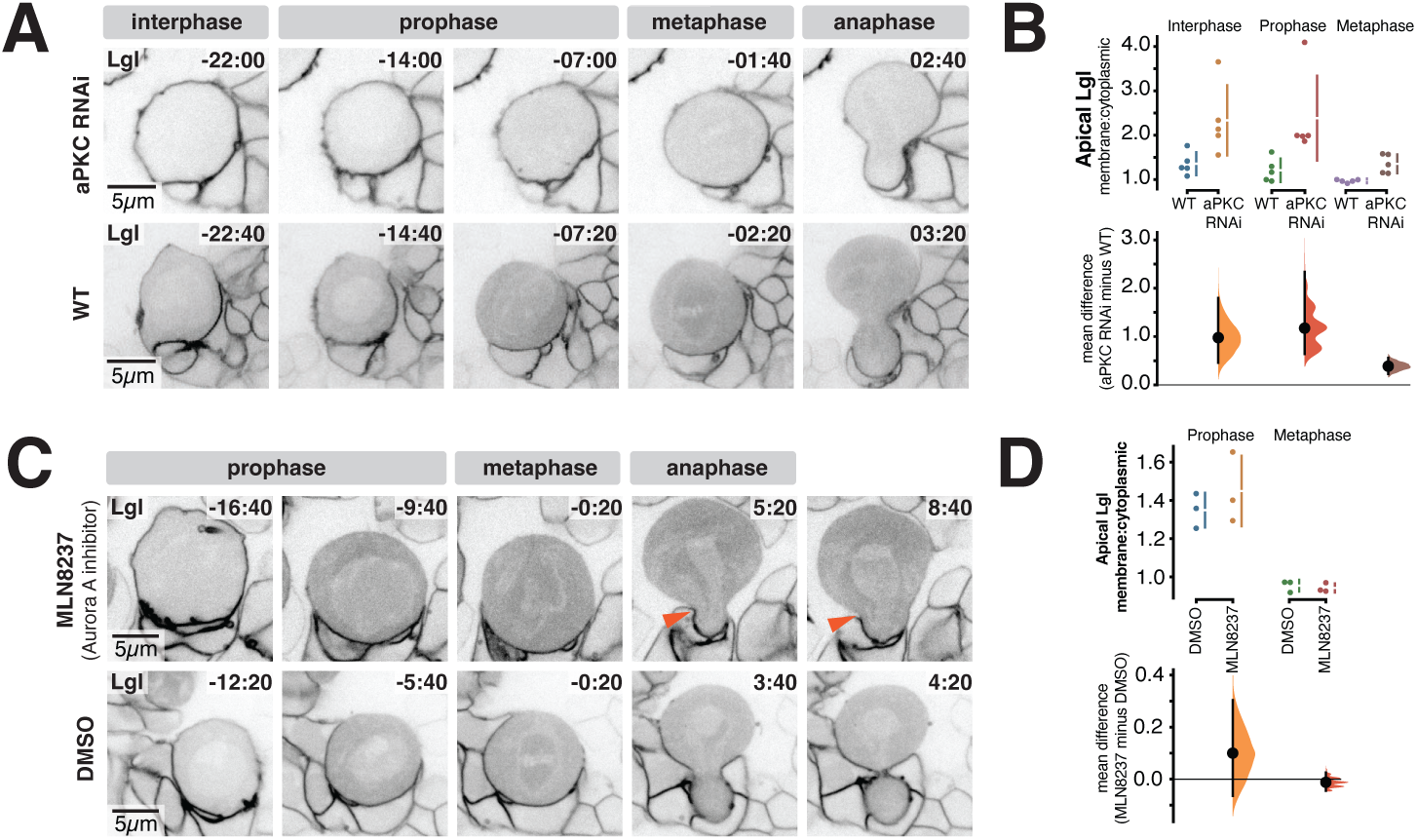
aPKC but not Aurora A is required for Lgl removal from the NSC membrane in early mitosis. (A) Lgl dynamics in an NSC from the intact central nervous system expressing aPKC RNAi. Frames from Video 2 are shown. The NSC is expressing Lgl-GFP and aPKC RNAi (both expressed from Worniu-GAL4 driven UAS). Time is shown relative to anaphase onset. The same timepoints are shown from an NSC expressing Lgl-GFP, but not expressing aPKC RNAi for a wild-type (”WT”) comparison. (B) aPKC is required for Lgl removal from the membrane in early mitosis. The Gardner-Altman estimation plot displays error bars indicating one standard deviation (with the gap representing the mean). The bar in the mean difference comparison reflects the bootstrap 95% confidence interval. Each data point corresponds to a measurement taken from a distinct NSC. (C) Lgl dynamics in an NSC from the intact central nervous system incubated with the specific Aurora A inhibitor MLN8237. Frames from Video 2 are shown. The NSC is expressing Lgl-GFP along with aPKC RNAi, both driven by Worniu-GAL4 and UAS. Arrowheads indicate furrow retraction during late cytokinesis, providing evidence of drug effectiveness. The time is shown relative to anaphase onset. For comparison, similar timepoints are depicted for an NSC incubated with DMSO alone (vehicle control). (D) Aurora A inhibition does not detectabley influence Lgl membrane displacement in early mitosis. Lgl apical membrane-to-cytoplasmic ration is shown for reated with either MLN8237 or the vehicle control (DMSO) during interphase, prophase, and metaphase. The Gardner-Altman estimation plot displays error bars representing one standard deviation (with the gap indicating the mean). The bar representing the mean difference comparison reflects the bootstrap 95% confidence interval.

We also tested if Aurora A activity is required for Lgl membrane displacement as it can directly phosphorylate Lgl and can also regulate aPKC polarity^5,7,20,25^. Consistent with our results indicating that aPKC is the dominant kinase responsible for displacement of Lgl from the membrane in NSCs, we found that inhibition of Aurora A activity with MLN8237 did not cause a significant difference in Lgl removal from the membrane compared to wild-type NSCs (Fig. 2C,D; Video 2). The patterned removal of Lgl from the membrane proceeded with similar dynamics and extent as observed in untreated cells. However, we did observe cytokinesis defects in treated NSCs (Fig. 2C; Video 2) indicating that Aurora A was successfully inhibited. We conclude that, despite its known ability to phosphorylate Lgl, Aurora A does not participate in the removal of Lgl from the NSC membrane. This conclusion is consistent with a previous study demonstrating that Aurora A phosphorylation sites on Lgl aren’t required for NSC membrane displacement^5^.

### Correlated aPKC and Lgl membrane dynamics

We next sought to determine the degree to which aPKC and Lgl membrane dynamics are spatially and temporally correlated. The recruitment of aPKC to the membrane in early prophase is a complex, multistep process where aPKC is initially localized to small domains on the apical membrane before coalescing into an apical cap by metaphase^26,27^. To determine the interplay between Lgl’s disappearance and aPKC’s appearance on the membrane, we simultaneously imaged aPKC-GFP (expressed from its endogenous promoter) and Lgl-mCherry (expressed by Worniu-GAL4 driven UAS) in NSCs. This approach allowed us to directly observe the spatial and temporal dynamics of aPKC recruitment with the pattern of Lgl removal from the cell membrane. Lgl removal at the apical pole of the NSC was tightly correlated with the appearance of aPKC at that site (Fig. 3A-C; Video 3). In the apical hemisphere, the appearance of aPKC at the membrane was strongly correlated with Lgl removal. However, Lgl was also removed from the basal membrane even though aPKC does not localize there. We conclude that aPKC removes Lgl from the membrane both directly (i.e. apical membrane) and indirectly (i.e. basal membrane). Indirect removal of basal Lgl by apical aPKC is likely influenced by factors such as the rate of Lgl diffusion along the plasma membrane.

**Figure 3.**
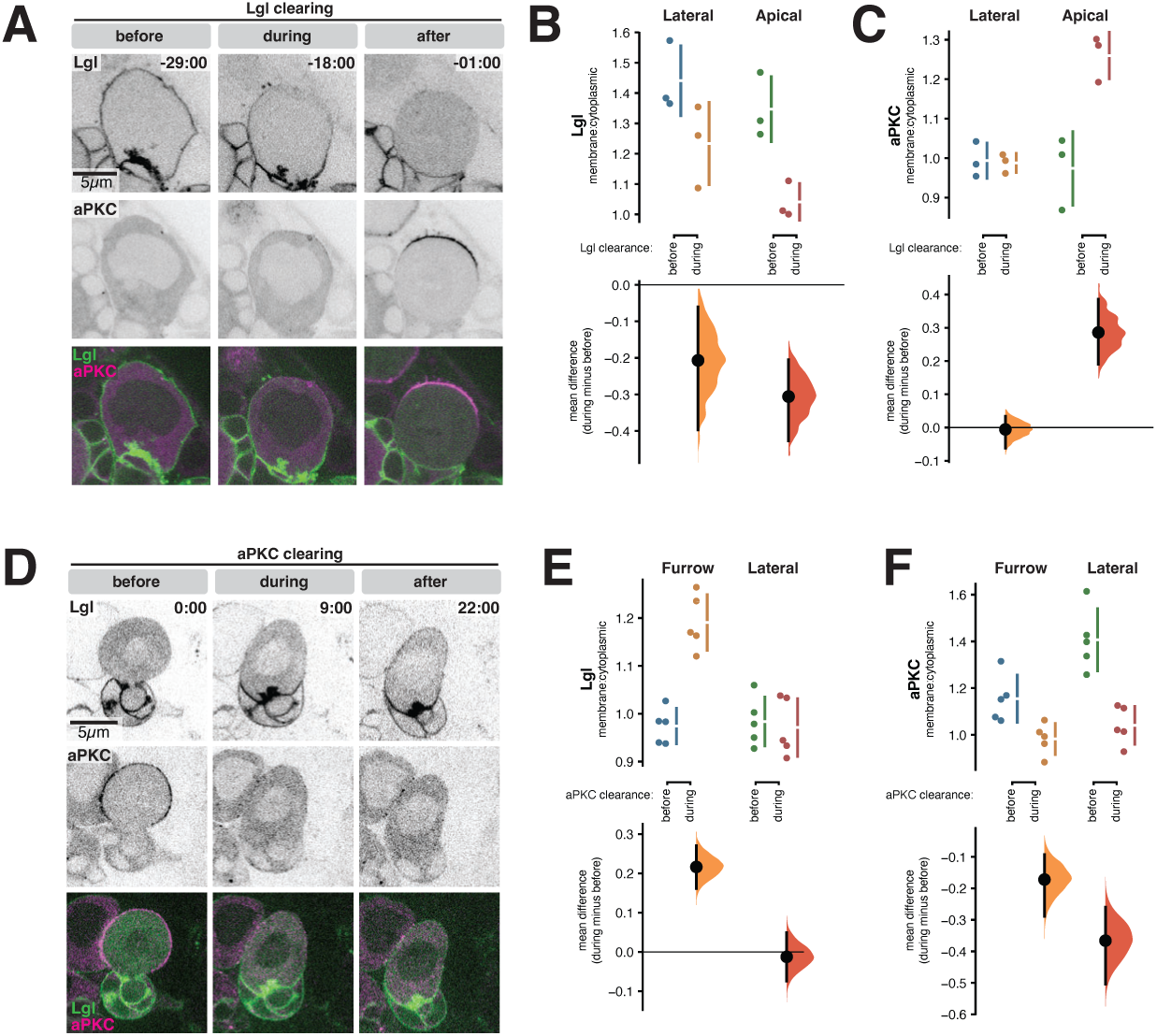
Correlated aPKC and Lgl NSC membrane dynamics. (A) Lgl and aPKC dynamics in an NSC from the intact central nervous system (CNS) during Lgl membrane clearance at mitotic entry. Frames from Video 3 are shown. The NSC is expressing aPKC-GFP (expressed from its native promoter) and Lgl-mCherry (expressed from Worniu-GAL4 driven UAS). Time is indicated relative to anaphase onset. (B) Lgl patterned removal at mitotic entry. The membrane-to-cytoplasmic ratio of Lgl was assessed both before and during the clearance of Lgl from the membrane, specifically at the apical and lateral membrane. The Gardner-Altman estimation plot displays error bars that indicate one standard deviation, with the gap denoting the mean value. The bar in the mean difference comparison illustrates the bootstrap 95% confidence interval. Each point on the plot corresponds to a measurement taken from a distinct NSC. Time is indicated relative to anaphase onset. (C) aPKC is apically enriched when Lgl begins clearing from the membrane at mitotic entry. The membrane-to-cytoplasmic ratio of aPKC is presented both before and during the clearance of Lgl from the membrane, measured at the apical and lateral membrane. Plotted as in (B) (D) The dynamics of Lgl and aPKC in an NSC from the intact CNS during the clearance of aPKC from the membrane at mitotic exit are illustrated with frames extracted from Video 3. The NSC is expressing aPKC-GFP driven by its native promoter and Lgl-mCherry expressed by Worniu-GAL4 driven UAS. Time is indicated relative to the beginning of the imaging session. (E) Lgl patterned membrane reloading at mitotic exit. At mitotic exit, as aPKC begins to clear from the membrane, Lgl is basally enriched at the site where the cytokinetic furrow reorganizes into the midbody. The ratio of membrane-to-cytoplasmic Lgl is presented both before and during the clearing of aPKC from the membrane, measured at the lateral membrane and near the furrow (as the midbody forms). The data is plotted as shown in panel (B). (F) aPKC patterned removal at mitotic exit. The ratio of membrane-to-cytoplasmic aPKC is presented both before and during the clearing of aPKC from the membrane, measured at the lateral membrane and near the furrow (as the midbody forms). The data is plotted as shown in panel (B)

Near the end of mitosis, aPKC spread along the nascent NSC membrane while being excluded from the membrane of the smaller nascent neural precursor (NP)^26^ (Figure 3D; Video 3). Shortly after it reached the furrow, aPKC was removed from the membrane. We observed patterned removal of aPKC from the membrane that was correlated with Lgl’s membrane reloading at mitotic exit (Figs. 1H,I, 3D-F; Video 3). As Lgl accumulated on the basal membrane of the nascent NSC near the cytokinetic pore, aPKC was removed from the membrane in a complementary pattern. Lgl continued to spread along the membrane until it occupied both basal and apical membrane and aPKC was no longer detectable on the membrane (Fig. 3D-F).

### Lgl excludes basal aPKC by regulating its interphase membrane association

The mutual exclusion model, which posits that Lgl and the Par complex reciprocally control each other’s localization by occupying opposing membrane domains, does not explain the behavior of these proteins in SOPs^20^ or NSCs^5^ (Fig. 3A). The primary inconsistency is the metaphase polarized state when the Par complex is polarized while Lgl is cytoplasmic. However, it is also not known whether Lgl is responsible for excluding aPKC from the membrane during interphase in NSCs. To gain insight into how Lgl regulates Par complex localization in these contexts, we examined aPKC localization in NSCs expressing Lgl RNAi (Worniu-GAL4 driven UAS). While aPKC is excluded from the cell membrane during interphase in wild-type NSCs, aPKC was not removed from the membrane at the end of mitosis in Lgl RNAi NSCs, remaining on the membrane throughout interphase and the subsequent mitosis (Fig. 4A-D; Video 4). However, the accumulation of aPKC at the apical membrane during early mitosis did not require Lgl, occurring both in wild-type and Lgl RNAi NSCs (Fig. 4A,C; Video 4). Apical enrichment caused aPKC to be polarized in both contexts at metaphase (Fig. 4E), with the key difference that Lgl RNAi NSCs had detectable basal aPKC at this stage (Fig. 4D). We conclude that Lgl is required to remove aPKC from the membrane at mitotic exit in NSCs but is not required for apical enrichment of the Par complex during early mitosis.

**Figure 4.**
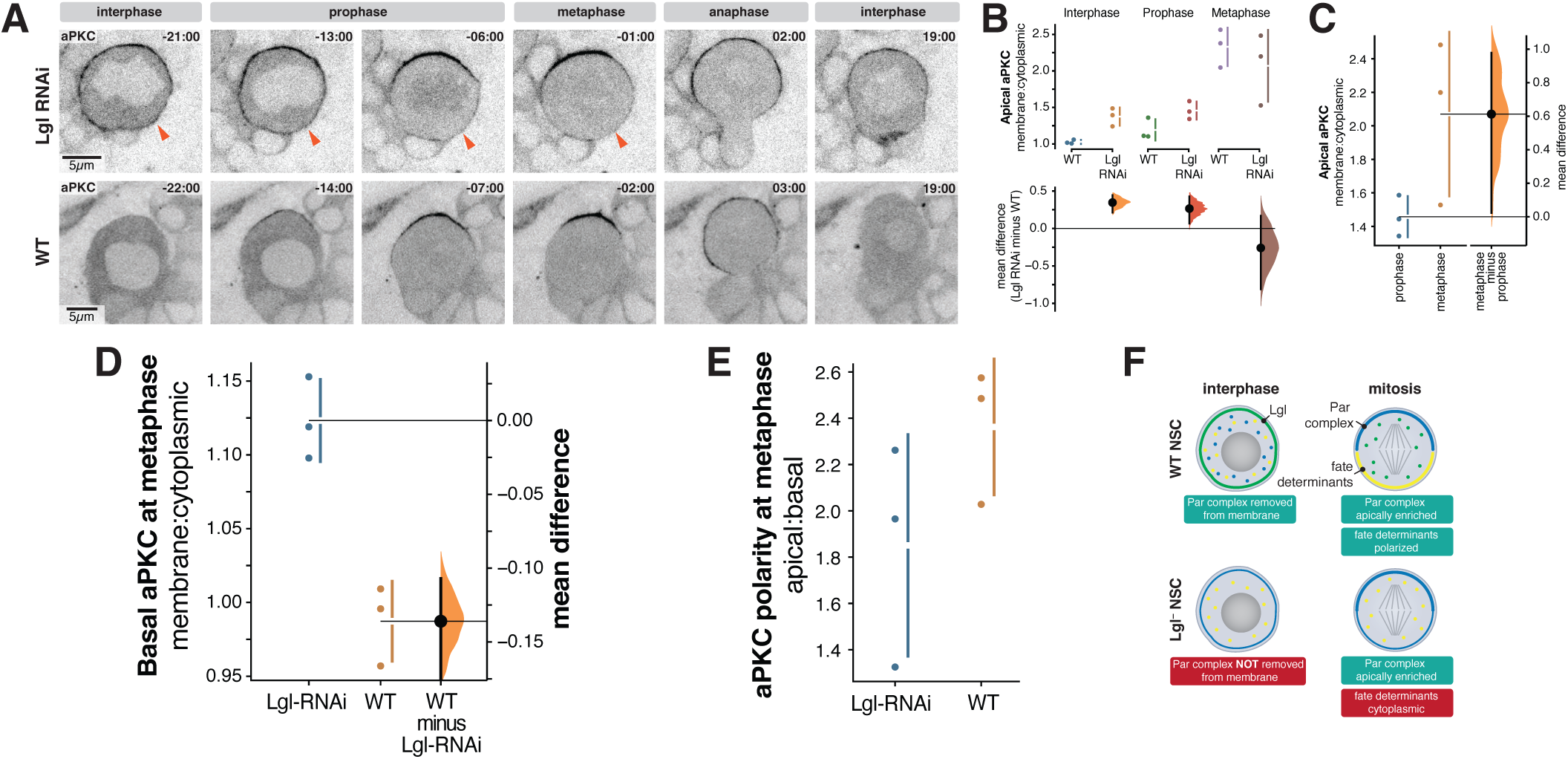
Lgl resets NSC polarity by removing aPKC from the membrane at mitotic exit. (A) aPKC dynamics in an NSC from the intact central nervous system expressing Lgl RNAi. The frames from Video 4 are presented, showcasing an NSC expressing aPKC-GFP, driven by its native promoter, and Lgl RNAi, induced by Worniu-GAL4 driving UAS. Arrowheads indicate the presence of basal aPKC before and during mitosis. Time is indicated relative to the onset of anaphase. For comparison, an NSC that does not express Lgl RNAi (wild-type "WT") is shown at corresponding time points. (B) Lgl is required to remove aPKC from the membrane during the return to interphase. A comparison of apical aPKC membrane enrichment in NSCs both expressing and not expressing Lgl RNAi during interphase, prophase, and metaphase stages. A Gardner-Altman estimation plot illustrates the data across three distinct NSC divisions. Error bars indicate one standard deviation, with the gap representing the mean. The bar in the mean difference comparison reflects the bootstrap 95% confidence interval. (C) aPKC is apically enriched at metaphase in Lgl RNAi NSCs. Comparing apical aPKC membrane enrichment at prophase and metaphase in Lgl RNAi expressing NSCs. Gardner-Altman estimation plot as in (B). (D) Lgl is required to remove aPKC from the basal membrane at metaphase. Comparison of basal aPKC membrane enrichment during metaphase in NSCs expressing Lgl RNAi versus those not expressing it. Gardner-Altman estimation plot as in (B). (E) Lgl is not required for metaphase aPKC polarity. The ratio of apical-to-cytoplasmic aPKC to basal-to-cytoplasmic aPKC during metaphase is depicted for three distinct NSC divisions. Error bars indicate one standard deviation, with the gap representing the mean value. (F) Polarity reset model for Lgl function in NSC asymmetric division. Lgl plays a crucial role in early interphase by removing the Par complex from the membrane, thereby establishing a cleared state essential for resetting polarity. The importance of clearing the Par complex from the membrane during each cell cycle is particularly evident in mitosis. Starting mitosis in a cleared state ensures that the Par complex binds exclusively to the apical membrane through its apical targeting mechanisms. As a result, the fate determinants can effectively polarize to the basal membrane without interference from the Par complex. Without Lgl, the Par complex persists on the membrane throughout interphase. During early mitosis, the Par complex accumulates at the apical membrane, similar to what occurs in wild-type NSCs, resulting in a polarized distribution. However, in this scenario, residual Par complex remains on the basal membrane during mitosis, effectively preventing fate determinants from binding to the basal membrane as they normally would.

### Lgl resets Par polarity for subsequent asymmetric NSC division

Our results provide a more nuanced view of Lgl function in Par polarity than provided by the mutual exclusion model. Rather than directly regulating metaphase Par polarity by occupying an opposing membrane domain, Lgl acts much earlier in the cell cycle to clear the Par complex from the NSC membrane at the end of mitosis. This mechanism ensures that each cell cycle begins without Par complex on the membrane (Fig. 4F). The importance of this “polarity reset” can be seen in NSCs where the Par complex isn’t removed from the membrane. In NSCs lacking Lgl function, the Par complex remains on the membrane after mitosis. Shortly after mitotic entry, the Par complex becomes enriched at the apical membrane, like in wild-type NSCs (Fig. 4C,E). However, residual basal Par complex from interphase (Fig. 4D) prevents fate determinant polarization^18^, disrupting asymmetric cell division. Thus, Lgl is not required for Par polarity in NSCs but instead ensures the Par complex is removed from the membrane when polarization begins in early mitosis. In other words, Par polarity is insufficient to support NSC asymmetric division – the Par complex must be on the apical membrane and not on the basal membrane. Basal Par complex, even at the reduced levels we observed compared to apically-enriched complex, disrupts fate determinant polarity and asymmetric cell division^18^.

While the metaphase polarized state has long been the focus of NSC asymmetric cell division research^2,28–31^, our results highlight the importance of the progression from mitosis to interphase to the process. Asymmetric cell division couples transitions in cell fate to the division cycle and fate transitions likely occur at or near mitotic exit. Consistent with the importance of this cell cycle phase, fate determinants are activated shortly after the end of mitosis^32^. The midbody that forms in the intercellular bridge connecting nascent siblings plays a key role in mediating asymmetric fate specification^32^. Intriguingly, Lgl membrane reloading and the concomitant displacement of aPKC into to the cytoplasm begins at or near the midbody (Fig. 3D-F) suggesting that fate specification and resetting polarity could be regulated by the same pathways.

The mechanism by which Lgl removes aPKC from the membrane at the end of mitosis remains an intriguing question. There are several possibilities based on our current understanding of Lgl and aPKC. One potential mechanism is that Lgl, upon its rapid return to the membrane post-mitosis, could act as a competitive inhibitor for aPKC membrane binding sites. Lgl’s strong affinity for the membrane^15,16^ might allow it to outcompete aPKC and displace it from the membrane. Alternatively, Lgl could recruit or activate a phosphatase that dephosphorylates aPKC or its binding partners, thereby reducing aPKC’s affinity for the membrane. Further investigation into these potential mechanisms will be crucial for fully understanding Lgl’s role in polarity regulation.

## Resource Availability

### Lead Contact

Contact the Lead Contact, Kenneth Prehoda, for further information or to request resources and reagents.

### Materials Availability

No new reagents were generated in this study.

### Data and Code Availability

Raw data available from the corresponding author on request.

## Experimental Model and Subject Details

### Fly Strains

Tissue-specific expression of UAS controlled transgenes in NSCs was achieved using a Worniu-GAL4 driver line^33^. Lgl dynamics were imaged using UAS-Lgl-GFP or UAS-Lgl-mCherry (as noted in figure and video legends). Plasma membrane dynamics were imaged using the membrane marker UAS-PLCδ-PH-mCherry, which expresses the pleckstrin homology domain of human PLCδ tagged with mCherry and binds to the plasma membrane lipid phosphoinositide PI(4,5)P_2_. Chromosome dynamics were monitored using the marker His2A-RFP. Imaging of aPKC was achieved using a BAC-encoded aPKC-GFP^34^. Worniu-GAL4 was used to express UAS-RNAi against aPKC and Lgl in NSCs. Knockdown was enhanced through the co-expression of UAS-Dicer2.

## Method Details

### Live Imaging

Third instar larvae were incubated at 29°C for at least 48 hours prior to imaging and dissection. Larvae were dissected in Schneider’s Insect Media (SIM) to obtain intact central nervous system explants, which were then mounted on a sterile poly-D-lysine coated 35mm glass bottom dish (ibidi Cat#81156) containing modified minimal hemolymph-like solution (HL3.1). Larval brain explants were imaged using a Nikon Eclipse Ti-2 Yokogawa CSU-W1 SoRa spinning disk microscope with dual Photometrics Prime BSI sCMOS cameras, utilizing a 60x water immersion objective. GFP-tagged proteins were visualized using 488 nm illumination, while RFP and mCherry tags were visualized using 561 nm illumination. Super-resolution images were captured via SoRa (super-resolution through optical photon reassignment) optics^35^. NSCs were identified based on size, location within the central nervous system, and expression of Worniu-GAL4 driven transgenes.

To acquire “dissociated NSCs”, *Drosophila* central nervous systems were collected from third instar larvae and incubated for 30 min at 30°C in Schneider’s Insect Medium containing 1 mg/mL Papain and 1 mg/mL Collagenase. After incubation, mechanical dissociation was performed by flushing medium/brains through a pipette tip. The resulting cell suspension was then pelleted using gentle centrifugation and washed 3 times with culture medium consisting of Schneider’s Insect Medium containing 10% synthetic fetal bovine serum, 20 mM Glutamine, 0.05 mg/mL Glutathione, and 0.01 mg/mL Insulin. After the final wash step, cells were resuspended and plated in culture medium consisting of Schneider’s Insect Medium containing 10% synthetic fetal bovine serum, 20 mM Glutamine, 0.05 mg/mL Glutathione, and 0.01 mg/mL Insulin and imaged with a Nikon Eclipse Ti-2 Yokogawa CSU-W1 SoRa spinning disk microscope with dual Photometrics Prime BSI sCMOS cameras, using a 60x water immersion objective.

### Pharmacological Inhibition

The selective Aurora A inhibitor MLN8237 (Alisertib) was solubilized in DMSO and used at a working concentration of 15 µm in all relevant experiments.

### Image Processing and Analysis

Imaging data was processed using ImageJ (FIJI package). To correct for photobleaching, the bleach correction tool was used.

Quantification for Fig. 1D: To quantify the dynamics underlying Lgl’s removal from the plasma membrane during early mitosis in NSCs within intact central nervous systems, we used ImageJ to acquire membrane:cytoplasmic ratios for Lgl-GFP signal intensity. To do this, we first measured Lgl membrane intensity by drawing a line with a 5 pixel width perpendicular to the membrane traversing from outside the NSC into the cytoplasm at two different locations, the apical pole and lateral membrane. Lgl membrane signal was quantified using the peak signal intensity along the line. Lgl cytoplasmic signal intensity was measured by taking the average signal intensity within a square ROI (region of interest) drawn inside the NSC cytoplasm, being careful to avoid sampling the nuclear compartment. These measurements allowed us to compare the apical membrane:cytoplasmic ratios to the lateral membrane:cytoplasmic ratios as NSCs moved through early mitosis. The figure depicts the lateral membrane:cytoplasmic ratio at the frame when the apical membrane:cytoplasmic ratio reached ∼1 (i.e. Lgl no longer detectable at the apical pole) for each NSC (n = 10).

Quantification for Figure 1G: To quantify the dynamics underlying Lgl’s removal from the basal plasma membrane, we measured the basal membrane:cytoplasmic ratio for Lgl-GFP signal intensity in dissociated NSCs. Dissociated NSCs provided a clearer view of the basal membrane compared to NSCs within intact central nervous systems, where the basal NSC membrane is obscured by the cluster of progeny cells in this region. The figure depicts the time relative to anaphase onset when the basal membrane:cytoplasmic ratio reached ∼1 (i.e. Lgl no longer detectable at the basal pole) for each NSC (n = 3). Anaphase onset was determined by using the chromosome marker His2a-RFP to identify the frame when chromosomes aligned at the metaphase plate began to separate.

Quantification for Figure 1J: To quantify the dynamics underlying Lgl’s rebinding to the plasma membrane during mitotic exit and interphase return, we measured membrane:cytoplasmic ratios for Lgl-GFP signal intensity in dissociated NSCs. Dissociated NSCs provided a clearer view of the basal membrane compared to NSCs within intact central nervous systems, where the basal NSC membrane is obscured by the cluster of progeny cells in this region. During mitotic exit and interphase return, cytokinesis is still underway and the nascent sibling cells remain connected by an intercellular bridge containing the cytokinetic pore^32^. Membrane:cytoplasmic ratios were measured at three different locations along the nascent NSC membrane, near the cytokinetic pore (the most basal point of the nascent NSC membrane), the lateral membrane of the nascent NSC, and at the apical pole of the nascent NSC. These measurements were performed at the end of anaphase (when the 2 sets of chromosomes had finished moving apart), early interphase, and later in interphase.

Quantification for Figure 2B: To quantify the effects of aPKC knockdown on Lgl’s membrane binding over the course of the cell cycle, apical membrane:cytoplasmic ratios for Lgl-GFP signal intensity were measured (as described in Fig. 1D). Ratios were measured at three different time points in the cell cycle. In all the NSCs we sampled, at 15 minutes before nuclear envelope breakdown (NEB) no NSC had begun the rounding up process that marks mitotic entry. We therefore used 15 minutes prior to NEB as the “interphase” time point. NSCs were in the process of rounding up at 7 minutes prior to NEB, and we therefore used this as the “prophase” time point. We used the frame before anaphase onset as the “metaphase” time point. NSCs expressing aPKC RNAi were compared to control cells not expressing RNAi (wild-type).

Quantification for Figure 2D: To quantify the effects of Aurora A inhibition on Lgl’s removal from the plasma membrane during early mitosis apical membrane:cytoplasmic ratios for Lgl-GFP signal intensity were measured (as described in Fig. 1D). “Prophase” measurements were taken at 6 minutes and 40 seconds prior to NEB. “Metaphase” measurement were taken at 20 seconds prior to anaphase onset. NSCs treated with the Aurora A inhibitor solubilized in DMSO were compared to control cells treated with DMSO alone.

Quantifications for Figure 3B and 3C: To determine if Lgl’s removal from the apical membrane is spatially and temporally correlated with the recruitment of aPKC to the apical membrane during mitotic entry, we used NSCs expressing Lgl-mCherry and aPKC-GFP (n = 3) to measure the lateral and apical membrane:cytoplasmic ratio for both aPKC and Lgl at the frame in which apical Lgl-mCherry signal reached the limit of detection (Lgl membrane:cytoplasmic ratio ∼ 1). Figure 3B depicts the lateral and apical membrane:cytoplasmic ratio of Lgl-mCherry signal intensity. Figure 3C depicts the lateral and apical membrane:cytoplasmic ratio of aPKC-GFP signal intensity.

Quantifications for Figure 3E and 3F: To determine if the removal of aPKC from the membrane is spatially and temporally correlated with the recruitment of Lgl to the membrane, we performed dual imaging of NSCs expressing Lgl-mCherry and aPKC-GFP (n = 5). The quantifications depict measurements taken during mitotic exit, at the frame in which aPKC-GFP signal near the cytokinetic pore of the nascent NSC reached the limit of detection (membrane:cytoplasmic ratio ∼1). Figure 3E compares the membrane:cytoplasmic ratio of Lgl-mCherry signal intensity at two different locations on the nascent NSC, at the basal membrane of the nascent NSC (near the cytokinetic pore) and the lateral membrane. Figure 3F compares the membrane:cytoplasmic ratio of aPKC-GFP signal intensity at two different locations on the nascent NSC, at the basal membrane of the nascent NSC (near the cytokinetic pore) and the lateral membrane.

Quantification for Figure 4B: To quantify the effects of Lgl knockdown on aPKC localization at the apical membrane, apical membrane:cytoplasmic ratios for aPKC-GFP signal intensity were measured at interphase (15 minutes prior to NEB), prophase (7 minutes prior to NEB), and metaphase (1 minute before anaphase onset). NSCs expressing Lgl-RNAi (n = 3) were compared to wild-type NSCs (n = 3).

Quantification for Figure 4C: To quantify the effects of Lgl knockdown on aPKC localization at the basal membrane, basal membrane:cytoplasmic ratios for aPKC-GFP signal intensity were measured at metaphase (1 minute before anaphase onset). NSCs expressing Lgl-RNAi (n = 3) were compared to wild-type NSCs (n = 3).

Quantification for Figure 4D: To quantify the effects of Lgl knockdown on the localization of aPKC to the basal membrane, the basal membrane:cytoplasmic ratio for aPKC-GFP signal intensity was measured at metaphase (1 minute before anaphase onset). NSCs expressing Lgl-RNAi (n = 3) and compared to wild-type NSCs (n = 3).

Quantification for Figure 4E: To quantify the effects of Lgl knockdown on aPKC polarity, aPKC signal intensity at the apical and basal poles were compared. The apical membrane:cytoplasmic and basal membrane:cytoplasmic ratio for aPKC-GFP signal intensity was measured at metaphase (1 minute before anaphase onset). To calculate the apical:basal ratio, the apical membrane:cytoplasmic ratio was divided by the basal membrane:cytoplasmic ratio. NSCs expressing Lgl-RNAi (n = 3) were compared to wild-type NSCs (n = 3).

## Statistical Analysis

Gardner-Altman estimation plots and 95% confidence intervals of datasets were prepared using the DABEST package^36^. Statistical details can be found in the relevant figure legends.

## Key Resources Table

| REAGENT or RESOURCE | SOURCE | IDENTIFIER |
| --- | --- | --- |
| **Experimental Model: Fly Strains** |  |  |
| aPKC-GFP | François Schweisguth Lab<sup>34</sup> |  |
| UAS-Lgl-GFP | Jürgen A. Knoblich Lab<sup>20</sup> |  |
| Histone H2a-RFP (His2a-RFP) (chromosome marker) | Bloomington *Drosophila* Stock Center (BDSC) | BDSC Cat# 23650<br>RRID: BDSC 23650 |
| Worniu-GAL4 (NSC driver line) | Chris Q. Doe Lab |  |
| aPKC-RNAi | Bloomington *Drosophila* Stock Center (BDSC) | BDSC Cat#67204<br>RRID: BDSC 67204 |
| Lgl-RNAi | Bloomington *Drosophila* Stock Center (BDSC) | BDSC Cat#38989<br>RRID: BDSC 38989 |
| UAS-Lgl-mCherry | Jürgen A. Knoblich Lab |  |
| UAS-PLCδ-PH-mCherry<br>(membrane marker) | Bloomington *Drosophila* Stock<br>Center (BDSC) | BDSC Cat#51658<br>RRID: BDSC 51658 |
| **Pharmacological Inhibitors** |  |  |
| MLN8237 (Alisertib) | Selleckchem | Cat#S1133 |

## Video Legends

### Video 1: Patterned Lgl membrane displacement and reloading during neural stem cell (NSC) divisions

*Part 1: Lgl dynamics in NSCs (imaged in intact central nervous systems).* Super-resolution videos of NSCs expressing Lgl-GFP (Lgl) and His2a-RFP (His2a). Time relative to anaphase onset is indicated. 3 medical section videos of dividing NSCs within intact central nervous systems are shown.

*Part 2: Lgl dynamics in dissociated NSCs.* Super-resolution videos of dissociated NSCs expressing Lgl-GFP (Lgl) and His2a-RFP (His2a). Time relative to anaphase onset is indicated. 3 medial section videos of dividing dissociated NSCs are shown.

*Part 3: 3D projections and medial sections of Lgl and membrane dynamics in NSCs.* Super-resolution videos of NSCs expressing Lgl-GFP (Lgl) and the membrane marker PLCδ-PH-mCherry (membrane). Time relative to nuclear envelope breakdown (NEB) is indicated. 3D projections and medial sections are shown for 3 dividing NSCs within intact central nervous systems.

### Video 2: The effects of aPKC knockdown and Aurora A inhibition on Lgl dynamics

*Part 1: The effects of aPKC knockdown on Lgl dynamics.* Super-resolution videos of NSCs expressing Lgl-GFP (Lgl) and aPKC RNAi. Time relative to anaphase onset is indicated. Medial sections are shown for 6 dividing NSCs within intact central nervous systems.

*Part 2: Lgl dynamics in NSCs (wild-type control).* Super-resolution videos of NSCs expressing Lgl-GFP (Lgl). Time relative to anaphase onset is indicated. Medial sections are shown for 6 dividing NSCs within intact central nervous systems.

*Part 3: Lgl dynamics in NSCs treated with Aurora A inhibitor.* Super-resolution videos of NSCs expressing Lgl-GFP (Lgl) and His2a-RFP (His2a) and treated with Aurora A inhibitor. Time relative to anaphase onset is indicated. Medial sections are shown for 3 dividing NSCs within intact central nervous systems.

*Part 4: Lgl dynamics in NSCs treated with DMSO.* Super-resolution videos of NSCs expressing Lgl-GFP (Lgl) and His2a-RFP (His2a) and treated with DMSO (vehicle control). Time relative to anaphase onset is indicated. Medial sections are shown for 3 dividing NSCs within intact central nervous systems.

### Video 3: Dual imaging of Lgl and aPKC dynamics in dividing NSCs

*Part 1: Lgl and aPKC dynamics during mitotic entry.* Super-resolution videos of NSCs expressing aPKC-GFP (aPKC) and Lgl-mCherry (Lgl). Time relative to anaphase onset is indicated. Medial sections are shown for 3 dividing NSCs within intact central nervous systems.

*Part 2: Lgl and aPKC dynamics during mitotic exit.* Super-resolution videos of NSCs expressing aPKC-GFP (aPKC) and Lgl-mCherry (Lgl). Time relative to the start of imaging is indicated. Medial sections are shown for 4 dividing NSCs within intact central nervous systems.

### Video 4: The effects of Lgl knockdown on aPKC dynamics

*Part 1: aPKC dynamics in NSCs expressing Lgl RNAi.* Super-resolution videos of NSCs expressing aPKC-GFP (aPKC) and Lgl-RNAi. Time relative to anaphase onset is indicated. Medial sections are shown for 3 dividing NSCs within intact central nervous systems.

*Part 2: aPKC dynamics in NSCs (wild-type control).* Super-resolution videos of NSCs expressing aPKC-GFP (aPKC). Time relative to anaphase onset is indicated. Medial sections are shown for 3 dividing NSCs within intact central nervous systems.

## Acknowledgments

We thank Eurico Morais-de-Sá for helpful discussions, Sarah Siegrist for a detailed dissociated NSC protocol, and Adam Fries for maintaining the microscope used in this study. This work was supported by NIH grants R35GM127092 and K99GM147601.

## Author Contributions

B.L., S.E.W., and K.E.P. designed the experiments. B.L. and S.E.W. performed the experiments. B.L., S.E.W., and K.E.P analyzed the data, prepared the figures, and wrote the manuscript.

